# Neural responses to binocular in-phase and anti-phase stimuli

**DOI:** 10.1101/2025.09.08.674974

**Authors:** Bruno Richard, Daniel H. Baker

## Abstract

Binocular vision fuses compatible inputs from the two eyes into a single percept, whereas incompatible inputs can produce rivalry, lustre, or diplopia. We measured neural responses to binocular stimuli with different phase relationships to test predictions from contemporary binocular combination models. Steady-State Visually Evoked Potentials (SSVEPs) were recorded from 15 observers in response to monocular and binocular stimulation at 3 Hz, using either On/Off or counterphase flicker with varied spatial and temporal phase relationships. On/Off flicker elicited responses at the fundamental frequency (3 Hz), and its integer harmonics, while counterphase flicker generated responses at the even integer harmonics (6Hz, 12Hz, 18Hz). Manipulating phase relationships modulated these response patterns, including a reduction in the fundamental amplitude for On/Off flicker. The data were modelled with a series of binocular combination algorithms, ranging in complexity from a simple linear sum to a two-stage binocular gain-control model with parallel monocular and binocular phase-selective channels. The model required parallel monocular channels to account for our data, whereas phase selectivity was not essential. Overall, the two-stage contrast gain-control model remains a powerful and flexible framework for describing binocular combinations across various experimental conditions and modalities.

## Introduction

The human visual system integrates input from both eyes to form a unified binocular representation of the world. Combining monocular inputs enhances sensitivity to the presented stimuli, particularly when the contrast is low or near detection threshold (Baker et al., 2018; Campbell and Green, 1965; Meese et al., 2006). Contrast sensitivity improves by at least 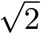 when stimuli are presented binocularly rather than monocularly (Baker et al., 2018; Blake and Wilson, 2011; Campbell and Green, 1965; Richard et al., 2018). Notably, the visual system also attempts to combine inputs even when the stimuli presented to each eye are markedly different (i.e., incompatible). In such cases, observers may perceive binocular rivalry (Blake, 1989; Wilson, 2003), diplopia, or visual lustre (Wendt and Faul, 2022). Despite differences in perceptual outcomes, computationally, the underlying processes of binocular combination for compatible and incompatible inputs appear similar (Baker et al., 2007a; Legge, 1984a). Psychophysical responses to compatible and incompatible stimuli can be effectively explained by a single psychophysical model that involves nonlinear transduction, followed by summation across monocular and binocular phase-selective channels (Baker et al., 2007a; Baker and Meese, 2007). Here, we explore if the integrative processes defined over multiple behavioural tasks are also reflected at the neural level.

It has long been known that stimuli presented binocularly are summed across the eyes. In contrast detection tasks, where stimulus contrast is low, binocular presentation increases sensitivity by approximately 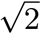 (Campbell and Green, 1965). This implies that observers require roughly 1.4 times more contrast to detect a monocular stimulus than a binocular one. The binocular improvement in sensitivity (Legge, 1984b) is consistent with a non-linearity operating before the signals from the two eyes (*c*_*L*_ and *c*_*R*_) are combined:

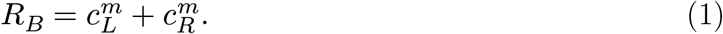

Here, the exponent *m* determines the degree of summation (*R*_*B*_). When *m* = 1, summation is linear, yielding a doubling of sensitivity. When *m* = 2, summation is reduced to 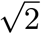. Several studies have reported summation ratios over 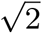 with some approaching 1.8 (Meese et al., 2006; Simmons, 2005; Simmons and Kingdom, 1998). A recent meta-analysis of 65 studies (N = 716) found an average binocular summation ratio of 1.5 (Baker et al., 2018). This work highlighted the challenges in accurately describing the binocular summation process (e.g., *m*) as individual variability and methodological differences can greatly impact binocular summation.

The contrast of stimuli is crucial in measuring binocular summation. Binocular summation can be very large when stimulus contrast is low; however, the binocular advantage is seldom observed in tasks that involve higher contrasts. In contrast discrimination tasks, where observers judge the contrast difference between otherwise identical stimuli, binocular presentation no longer confers a benefit in sensitivity: discrimination thresholds are identical whether stimuli are shown to one eye or both (Legge, 1984a; Maehara and Goryo, 2005; Meese et al., 2006). However, this does not indicate that binocular summation does not occur at higher contrasts; the monocular signals are still summed (Meese et al., 2006; Meese and Baker, 2011). Instead, the advantage of summation is counteracted by normalization mechanisms that maintain consistency across viewing conditions (referred to as ‘ocularity invariance’). In gain control models of early vision, normalization can stem from interocular and self-suppressive signals (Meese et al., 2006):

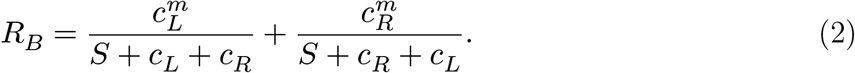

When both eyes are stimulated, the suppressive terms on the denominators offset the enhanced excitatory signals, nullifying the binocular advantage. The new parameter, S, is included to set overall sensitivity of the monocular response.

A similar pattern of results is observed in neural measurements of binocular summation. In functional Magnetic Resonance Imaging (fMRI) recordings, neural responses are significantly larger for binocular than monocular responses when stimulus contrast is low (Moradi and Heeger, 2009). At higher contrasts, observers no longer show the binocular advantage. As with behavioural results, these findings are well-explained computationally by interocular suppression and binocular contrast normalization. The equivalency in binocular summation between psychophysics and neuroimaging findings means that both data types can constrain binocular summation models. We, for example, have demonstrated that a popular model of binocular summation could be easily adapted to capture Steady-State Visually Evoked Potentials (SSVEPs) to monocular and binocular stimuli when one eye was occluded by a neutral density filter (Richard et al., 2018). Placing a neutral density filter in front of one eye darkens its input, reducing the amplitude and altering the phase of SSVEPs to stimuli presented to the filtered eye. Steady-State response amplitudes and phases to stimuli presented through different neutral density filter strengths were well described by a model that first used a biophysically plausible temporal filter on the input stimuli, followed by self and interocular suppression, and finally, binocular contrast normalization. This model also explained psychophysically measured binocular summation in the same group of observers. A comprehensive description of the processes involved in binocular summation should explain behavioural and neuroimaging findings under various experimental conditions of binocular summation.

Many studies have worked towards the development of a comprehensive description of the process of binocular summation in human vision using a variety of psychophysical and neuroimaging approaches (Baker et al., 2008, 2007b; Campbell and Green, 1965; Ding et al., 2013; Ding and Sperling, 2006; Legge, 1984a; Lygo et al., 2021; Maehara and Goryo, 2005; May and Zhaoping, 2016; Richard et al., 2018). This is not an insignificant challenge; such a description must account for multiple components of early vision, including identifying relevant signals, defining how these signals might interact, what non-linearities are present, and most importantly, how these signals are summed and/or differenced. Early approaches (Campbell and Green, 1965) proposed the 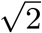 improvement from binocular summation could be explained because the visual system integrates signal and uncorrelated noise from the monocular channels. However, it was also recognized (Blake et al., 1981b; Campbell and Green, 1965) that the model can only return 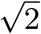 improvement when the noise in one eye is ignored, or that the observer is able to respond selectively to the monocular pathway in monocular trials. One model that has proven very informative and capable of describing binocular summation under various experimental conditions is the two-stage contrast gain control model developed by Meese et al. (2006). In a two-stage process, the model captured detection and discrimination thresholds for monocular and binocular presentation, in addition to dichoptic masking (when the stimuli presented to both eyes are not identical). First, the input is rectified by an excitatory non-linearity (*m* ≈ 1.3) and normalized by self and interocular suppression (see Equation 2). The binocular (e.g., combined) signal then undergoes a second contrast normalization before the decision stage,

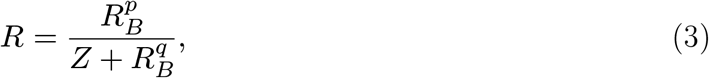

where the exponents *p* and *q* typically have quite large values, and determine the shape of the contrast discrimination (‘dipper’) functions. As with the parameter S in Equation 2, the parameter Z is included to fix the overall sensitivity of the binocular response.

Subsequent iterations of the two-stage contrast gain control model added channels for opposite contrast polarities (Baker and Meese, 2007), and monocular channels parallel to the binocular summing channel (Georgeson et al., 2016). Polarity-specific channels were included to explain masking effects when the stimuli presented to each eye had opposite phase polarities (i.e., dichoptic presentation). Antiphase stimuli do not cancel each other out (as would be expected if their luminances were linearly summed): the stimuli remain detectable and the two inputs can sum (Bacon, 1976; Baker and Meese, 2007; Simmons, 2005), at least in a probabilistic sense. Parallel monocular channels are consistent with adaptation after-effects that suggest monocular signals may be preserved and available for perception following binocular summation (Blake et al., 1981a; Moulden, 1980).

While the possibility of monocular channels had been considered (Legge, 1984a), many assumed that only the binocularly summed signal contributed to perception. Georgeson et al. (2016) developed a specific experimental condition to assess the involvement of monocular channels. They devised a discrimination task where the target interval presented a contrast increment to one eye (e.g., 10% pedestal + 2%) and a contrast decrement to the other (e.g., 10% pedestal - 2%). If the only available signal is a binocularly summed one, the task would be nearly impossible to complete; the target interval would be perceptually identical to the pedestal-only interval. However, observers were able to complete the task. The two-stage contrast gain control model with parallel monocular channels was the only model able to capture observer thresholds from all experimental conditions, including the binocular increment and decrement tasks. Evidence of an additional differencing channel accompanying the summing channel commonly described in work on binocular combination (Chen and Li, 1998; Li and Atick, 1994; May et al., 2012; May and Zhaoping, 2016) has also accumulated. As the name implies, this channel encodes the difference between stimuli presented to the left and right eye; it is involved in early contributions to disparity processing and stereovision. While computational models of binocular vision do not explicitly include differencing channels, encoding the difference between monocular signals is, indirectly, included in the two-stage contrast gain control model defined by Georgeson et al. (2016).

Steady-State Visually Evoked Potentials offer a unique opportunity to bridge the gap between models developed from psychophysical data and neuroimaging, as responses to stimulus contrast (i.e., SSVEP amplitude) are directly associated with behavioural sensitivity to contrast (Norcia et al., 2015; Wade and Baker, 2025). Two common stimulus presentation protocols are used to generate SSVEPs: a sinusoidal On/Off flicker, where the stimulus alternates between a blank background (0% contrast) and the peak contrast, and sinusoidal counterphase flicker, where the stimulus alternates in phase (i.e., the black regions become white, and the white regions become black). On/Off flicker activates different populations of on and off-cells independently once per cycle, generating frequency-following responses at the frequency of the sinusoidal modulation (1F). On/Off flicker can also generate SSVEPs at the odd and even integer harmonics (2F, 3F, 4F, etc.), which reflect nonlinear processing in the visual system (Regan and Regan, 1988). Counterphase flicker will generate two transients per cycle, resulting in SSVEPs at the even harmonics of the flicker frequency (2F, 4F, 6F, etc.). These frequency-doubling responses are argued to provide a cleaner measure of nonlinear visual responses (Kim et al., 2011; Skottun and Skoyles, 2007). Using both stimulation protocols yields the data needed for a comprehensive description of linear and nonlinear visual processes, thereby enabling the evaluation of psychophysical models of visual perception.

The current architecture of the two-stage contrast gain control model has been rigorously evaluated on psychophysical data, providing a solid foundation for our understanding of binocular combination (Baker and Meese, 2007; Georgeson et al., 2016; Meese et al., 2006). While previous studies have applied the two-stage contrast gain control model to neuroimaging data (Lygo et al., 2021; Richard et al., 2018), they only included experimental conditions where the phases of the sinusoidal gratings presented to each eye were identical. To accurately assess the current architecture of the two-stage contrast gain control model, neuroimaging data for binocular presentation of stimuli in opposite phase polarity are required. Here, we recorded SSVEPs to monocular and binocular stimuli with different spatial and temporal phase relationships and stimulation protocols (On/Off and counterphase flicker) to obtain the data needed to evaluate the two-stage contrast gain control model. By progressively increasing the complexity of the model, we demonstrate that many components of binocular combination, such as monocular non-linearities, interocular interactions, and parallel monocular channels, are required to explain neural responses to our set of experimental conditions. The two-stage contrast gain control model remains a powerful and flexible descriptor of the architecture of binocular combination for data collected across many experimental conditions and modalities.

## Methods

### Participants

Fifteen observers (*N*_*male*_ = 4; age range [19 34]), including authors BR and DHB, participated in this study. All observers had normal or corrected-to-normal visual acuity, as verified by a Snellen chart, and normal binocular vision, as verified by a Titmus test. Written informed consent was obtained from all participants, the experimental procedures followed the Declaration of Helsinki guidelines, and the study was approved by the ethics committee of the Department of Psychology at the University of York.

### Apparatus

All stimuli were presented using a gamma-corrected ViewPixx 3D display (VPixx Technologies, Canada) driven by a Mac Pro. Binocular separation with minimal crosstalk was achieved by synchronizing the display’s refresh rate with the toggling of a pair of Nvidia stereo shutter goggles using an infrared signal. The monitor refresh rate was set to 120 Hz; each eye updated at 60 Hz (every 16.67 msec). The display resolution was set to 1920 × 1080 pixels. A single pixel subtended 0.027° of visual angle (1.63 arc min) when viewed from 57 cm. The mean luminance of the display viewed through the shutter goggles was 26 cd/m^2^. EEG signals were recorded from 64 electrodes distributed across the scalp according to the 10/20 EEG system (Chatrian et al., 1985) in a WaveGuard cap (ANT Neuro, Netherlands). We monitored eye blinks with an electrooculogram consisting of bipolar electrodes placed above the eyebrow and the cheek on the left side of the participant’s face. Stimulus-contingent triggers were sent from the ViewPixx display to the amplifier using a parallel cable. Signals were amplified and digitized using a PC with the ASAlab software (ANT Neuro, Netherlands). All EEG data were imported into MATLAB (Mathworks, MA, USA) using components of the *EEGlab* toolbox (Delorme and Makeig, 2004) and then exported for subsequent offline analysis using R.

### Stimulus Creation

Observer SSVEPs were measured with a single horizontal sinusoidal grating that sub-tended 15° of visual angle on the retina with a spatial frequency of 3 cycles/° of visual angle (Figure 1A). All stimuli were generated using *MATLAB* (MathWorks, Natick, MA) and presented to observers using *Psychtoolbox* (Brainard, 1997; Kleiner et al., 2007; Pelli, 1997). Our experimental conditions modulated the interocular spatial phase of stimuli (Figure 1A). Under binocular viewing, the sinusoidal gratings could be presented in spatial phase or spatial anti-phase. When stimuli were presented in spatial phase, the aligned sinusoidal gratings were identical in both eyes (Δ*ϕ* = 0). The spatial anti-phase condition phase-shifted one of the sinusoidal gratings by 180° (Δ*ϕ* = *π*). Stimuli were also modulated in their oscillatory pattern, which could be On/Off or counterphase flicker at a frequency of 3Hz (Figure 1B). Under On/Off contrast flicker, the relative contrast of the gratings began at 0%, increased smoothly to 100% of the nominal maximum (100% Michelson contrast), and then returned to 0% over 333 ms (i.e., one cycle). On/Off flicker will generate SSVEPs at the fundamental frequency (3Hz) and its integer harmonics (2F, 3F, 4F, see Figure 1C). Counterphase flicker reversed the phase of the gratings at a frequency of 3Hz. The contrast of the grating began at the relative maximum (100%), gradually decreased to 0% of the relative maximum, and then increased again to 100% but in the opposite phase polarity. Unlike On/Off flicker, counterphase flicker generates two nearly identical transients per cycle and thus does not produce SSVEPs at the fundamental frequency (3Hz) but only its even harmonics (Wade and Baker, 2025).

**Figure 1:**
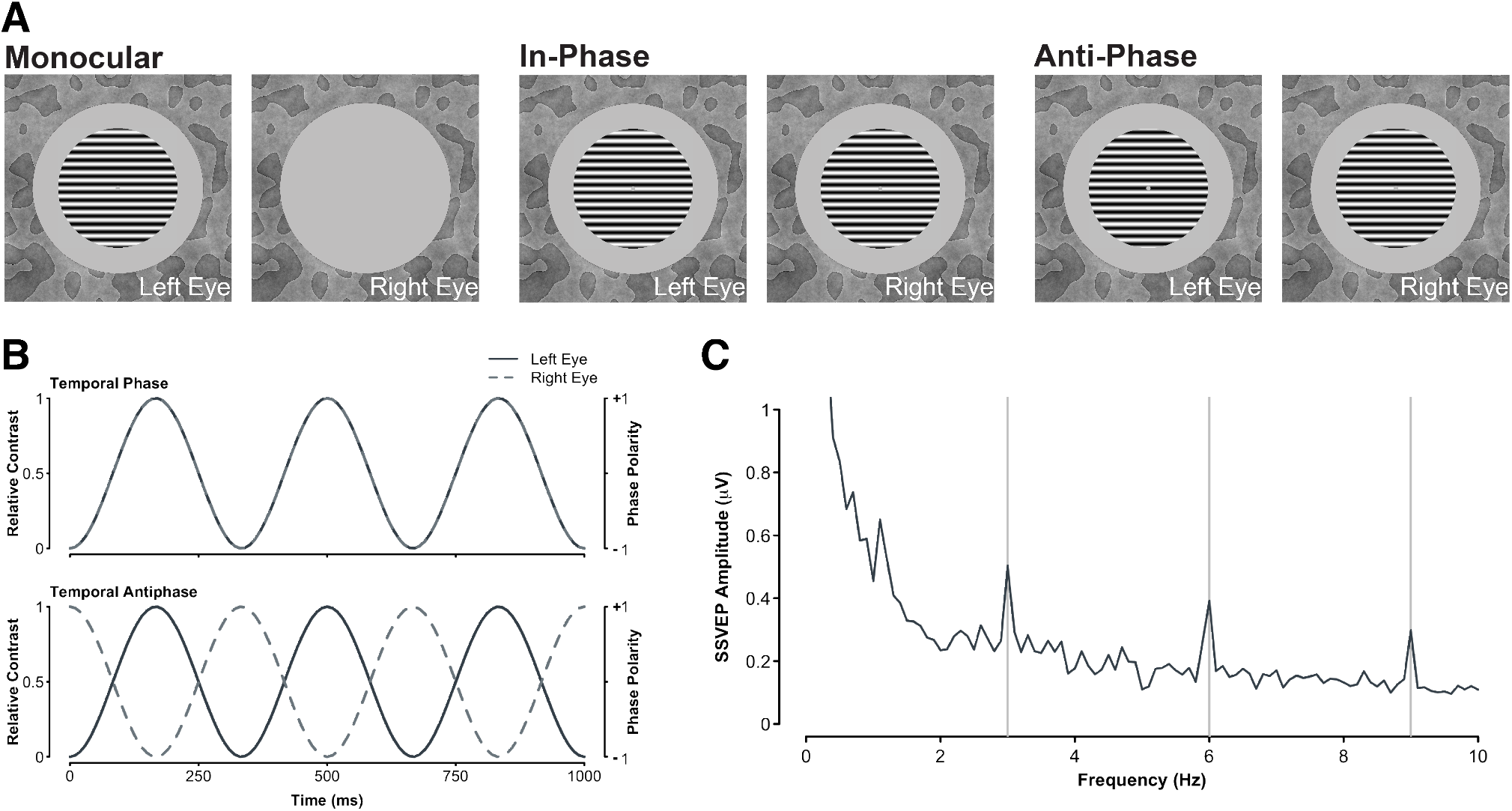
**A**. The spatial configuration of stimuli presented to observers in our experiment. Monocular conditions presented the sinusoidal grating to the left or right eye of observers (counterbalanced) while the other eye was presented with a gray screen set to mean luminance. Binocular conditions could be shown with stimuli in spatial phase, whereby the phase of both sinusoidal gratings was identical, or in spatial anti-phase, where the phase of the sinusoidal gratings presented to each eye was opposite. The background texture did not change throughout the trial to aid with binocular fusion. **B**. The temporal configuration of our stimuli. To generate SSVEPs, stimuli were contrast modulated in two ways: on/off (left Y axis) or counterphase (right Y axis). The oscillatory pattern could also be in phase (upper plot), where both stimuli were modulated in the same manner, or in counterphase (lower plot), where as one stimulus increased in contrast, the other decreased in contrast (or increased in opposite polarity). **C**. Example SSVEPs generated under binocular spatial and temporal in-phase viewing of stimuli for one observer, averaged across four electrodes (*Oz, POz, O1, O2*) and 12 repetitions.

To aid with binocular fusion, stimuli were surrounded by a static binocular texture presented beyond the central 19° stimulus aperture. These textures were constructed by first low-pass filtering a white (amplitude ∝ 1/*f* ^0^) noise pattern, dichotomizing its output into a binary image, and taking its phase spectrum. A second flat (amplitude ∝ 1/*f* ^0^) was adjusted by multiplying each spatial frequency’s amplitude coefficient by *f*^−1^ to generate a pink amplitude spectrum (Hansen and Hess, 2006; Tadmor and Tolhurst, 1994). The pink amplitude spectrum and the phase spectrum of the binary image were rendered in the spatial domain by taking the inverse Fourier transform, resulting in the pattern shown in Figure 1A. The surrounding texture was fixed during a trial, but its phase structure was changed across trials.

### Procedures

Steady-State Visually Evoked Potentials (SSVEPs) were recorded with monocular and binocular stimulation using either on-off or counterphase flicker at 3Hz. Across the eyes, binocular stimuli could be in spatial and temporal phases, temporal phase but spatial antiphase, spatial phase but temporal anti-phase, or spatial and temporal anti-phase (on-off flicker only). Stimuli presented in spatial and temporal anti-phase under counterphase flicker are identical to stimuli presented in spatial and temporal phase (and so we did not duplicate this condition). Thus, this experiment was comprised of nine conditions - two monocular and seven binocular - each repeated 12 times for a total of 108 trials. Stimulus presentation was separated into four experimental blocks, each containing 27 trials. A trial lasted 15 seconds; a grating stimulus flickered onscreen for 11 seconds, followed by a screen with its central 19° set to mean luminance for 4 seconds. Participants completed all 27 trials of an experimental block in a single sequence (6.75 minutes) and were given breaks between experimental blocks. The trial order was pseudo-randomized on each block. Participants did not receive explicit task instructions other than to fixate the marker in the center of the display and blink only during the blank period between stimulus presentations.

### SSVEP Analysis

We used whole-head average referencing to normalize each electrode to the mean signal of all 64 electrodes (for each sample point). The EEG waveforms were Fourier transformed at each electrode for a 10-second window, beginning one second after the stimulus onset to avoid onset transients. The Fourier spectra were coherently averaged (i.e., retaining the phase information) across four occipital electrodes (*Oz, POz, O1*, and *O2*) and trial repetitions (see Figure 1C). We then calculated signal-to-noise ratios (SNRs) by dividing the absolute amplitude in the signal bin (e.g., 3 Hz) by the mean of the absolute value of the ten adjacent bins (±0.5 Hz in steps of 0.1 Hz). There were no differences in the amplitude of the adjacent, noise, bins across experimental conditions. Given that distributions of ratios (including SNRs) are inherently skewed, the median SNR was taken across all participants; the median is a more robust descriptor of central tendency in skewed distributions.

## Results

Figure 2 shows the cross-participant median SNR spectra for all experimental conditions. Responses for all On/Off flicker experimental conditions generated peaks at the fundamental frequency (3Hz) and its harmonics (integer multiples of 3Hz). Similarly, counterphase flicker produced responses at twice the flicker frequency (6 Hz) and its harmonics. We assessed differences in SNR magnitude across experimental conditions via a permutation test, allowing a non-parametric comparison of a statistic between two conditions. We first take the median difference between two experimental conditions (i.e., the observed difference) to conduct the permutation tests. A null distribution is then constructed by combining SNR values from both experimental conditions, and randomly sampling them without replacement to create two groups of sizes identical to their original, but with values not associated with a particular experimental condition. The median difference of the randomly sampled SNRs is then taken. This process is repeated multiple times (e.g., *N* = 10 iterations) to build a distribution of median differences with no association of experimental condition (i.e., a null-hypothesis distribution). The observed median difference is then compared to this distribution. The proportion of scores greater than the observed difference represents the *p* value associated with the test. When comparing SNRs at the fundamental frequency (3Hz) for On/Off flicker, we find no statistically significant difference in median SNR magnitude between experimental conditions where stimuli were presented in temporal phase (see Figure 3). Monocular and binocular presentation for stimuli presented in temporal phase resulted in a similar response pattern under both On/Off and counterphase flicker modulations. This is consistent with ocularity invariance; binocular and monocular stimuli appear equal in magnitude at high contrast (Baker et al., 2007a; Meese et al., 2006).

**Figure 2:**
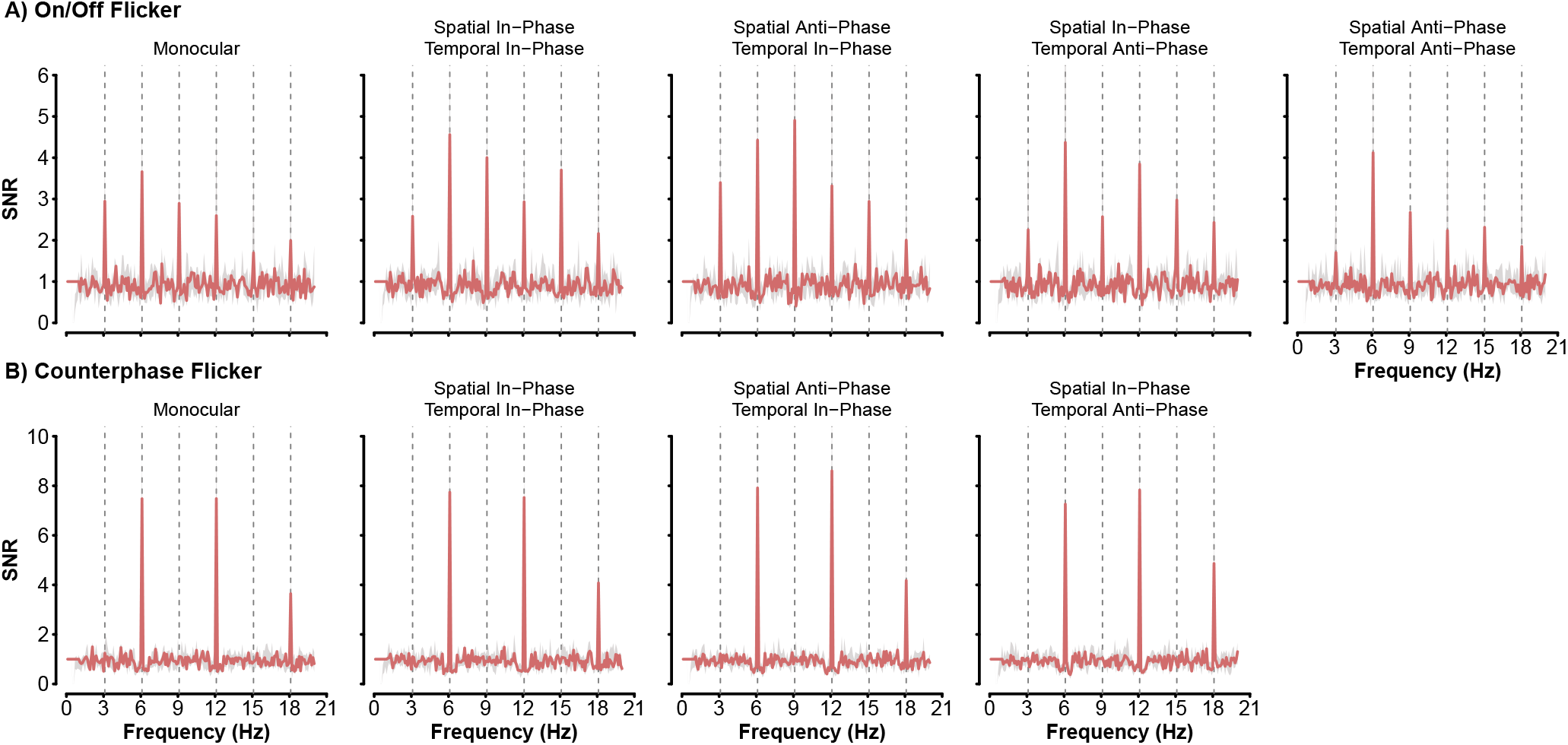
Cross-participant median SNRs for frequencies up to 20Hz. SNRs generated by On/Off flicker are shown in the top row (A), while those generated by counterphase flicker are shown in the bottom row (B). The light gray area represents bootstrapped 95% confidence intervals that were calculated by resampling (with replacement) participant SNRs 2000 times.

**Figure 3:**
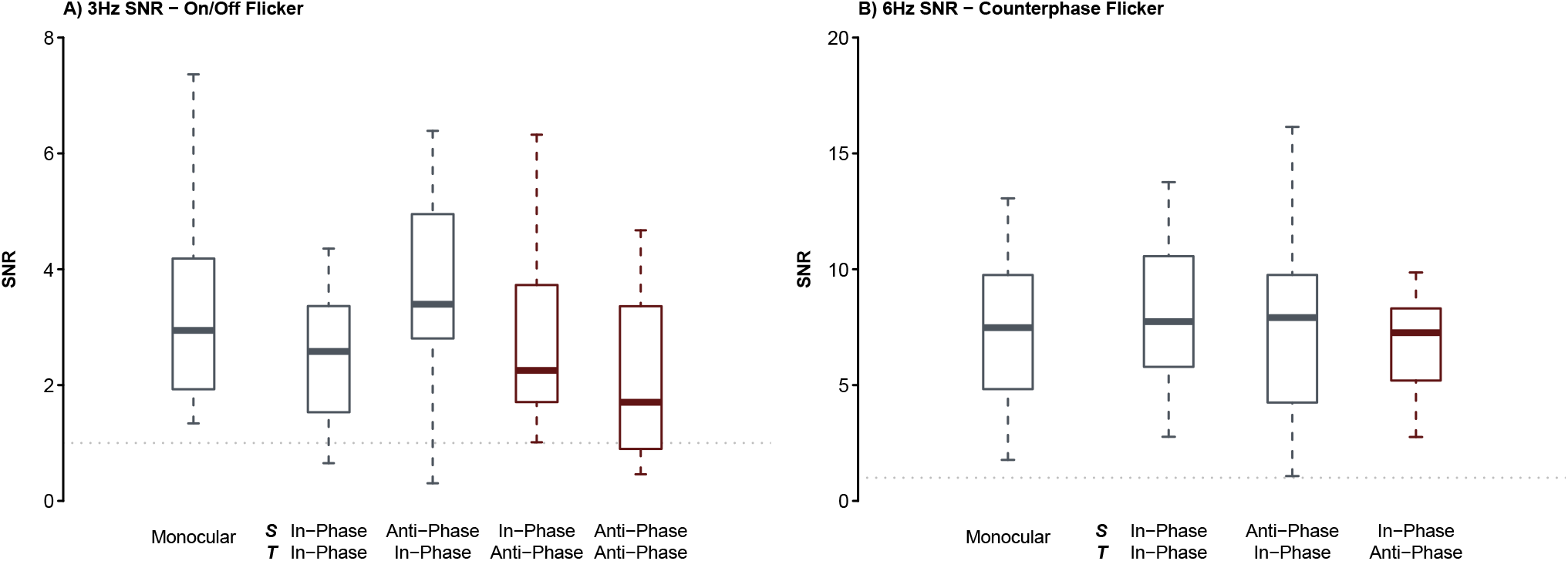
Boxplots of participant SNRs at the fundamental frequency (3Hz) under On/Off stimulation and its second harmonic for counterphase stimulation. A) Boxplots represent the distribution of participant SNRs at 3Hz. The median SNR is shown by the thick line within the box, with the lower and upper border of the box representing the first (25%) and third (75%) quartile of the distribution. The lower and upper whisker limits represent 1.5 times the interquartile range (distance between the third and first quartile). Boxplots for the binocular conditions have labels for their spatial (S) and temporal (T) phase relationships. Experimental conditions where stimuli are presented in temporal anti-phase are shown in maroon. B) As in A, boxplots show the distribution of participant SNRs at 6Hz. In both graphs, the dashed line represents an SNR of 1.0.

Changing the phase relationships of stimuli under On/Off flicker had some interesting impacts on the fundamental frequency (Figure 3A). Stimuli presented in spatial phase and temporal anti-phase generated smaller SNRs (median_SNR_ = 2.25) than stimuli presented in spatial anti-phase and temporal phase (median_SNR_ = 3.4, *p* =.047). The reduction in the amplitude relative to stimuli in spatial anti-phase and temporal phase was also observed for stimuli in temporal and spatial anti-phase (median_SNR_ = 1.7; *p* =.023). No other statistically significant difference in median SNRs were observed for all counterphase flicker conditions or the other harmonics for On/Off flicker (all *p* values were greater than.05). The median SNRs of temporal anti-phase stimuli under On/Off flicker, while smaller than those of other conditions, were nevertheless statistically significantly greater than 1.0 (spatial in-phase: *p* <.001 and spatial anti-phase: *p* =.004). The presence of a 3Hz response for binocular stimuli presented in temporal anti-phase is a strong indication that monocular responses remain and contribute to the SSVEP, as these conditions generate two transients per cycle. A purely binocular signal would only generate responses at 6Hz (Blake et al., 1981a; Georgeson et al., 2016; Moulden, 1980).

### Modelling

The perception of binocular stimulus contrast is well-explained by psychophysical models that process input contrast in two sequential contrast gain control stages interposed by binocular summation (Baker et al., 2008, 2007a, 2007b; Baker and Meese, 2007; Meese et al., 2006). This simple, yet powerful, family of models not only captures behavioural data well, but can also explain neural responses to binocular and dichoptic stimuli (Baker and Wade, 2017; Lygo et al., 2021; Richard et al., 2018). While other computational descriptions of binocular combination have been proposed (Blake and Wilson, 2011; Campbell and Green, 1965; Ding and Sperling, 2006), the goal of this manuscript is to probe the architecture of the two-stage contrast gain control model and its subsequent variants with a rich SSVEP data set. Our SSVEP results show the expected pattern of binocular combination for stimuli presented at high contrast (i.e., ocularity invariance) but also intriguing effects that are likely explainable by the most recent extension of the two-stage contrast gain control model, as defined by Georgeson et al. (2016). Our goal in this modelling section is not to argue for a novel description of binocular summation but to explore the architecture required under the two-stage contrast gain control framework, to describe our effects. Thus our approach is to progressively increase the complexity of the binocular combination model, beginning with a deliberately wrong model (i.e., linear combination) and building up to a multi-channel model (Georgeson et al., 2016) with monocular, binocular, and phase-selective pathways (Figure 4).

**Figure 4:**
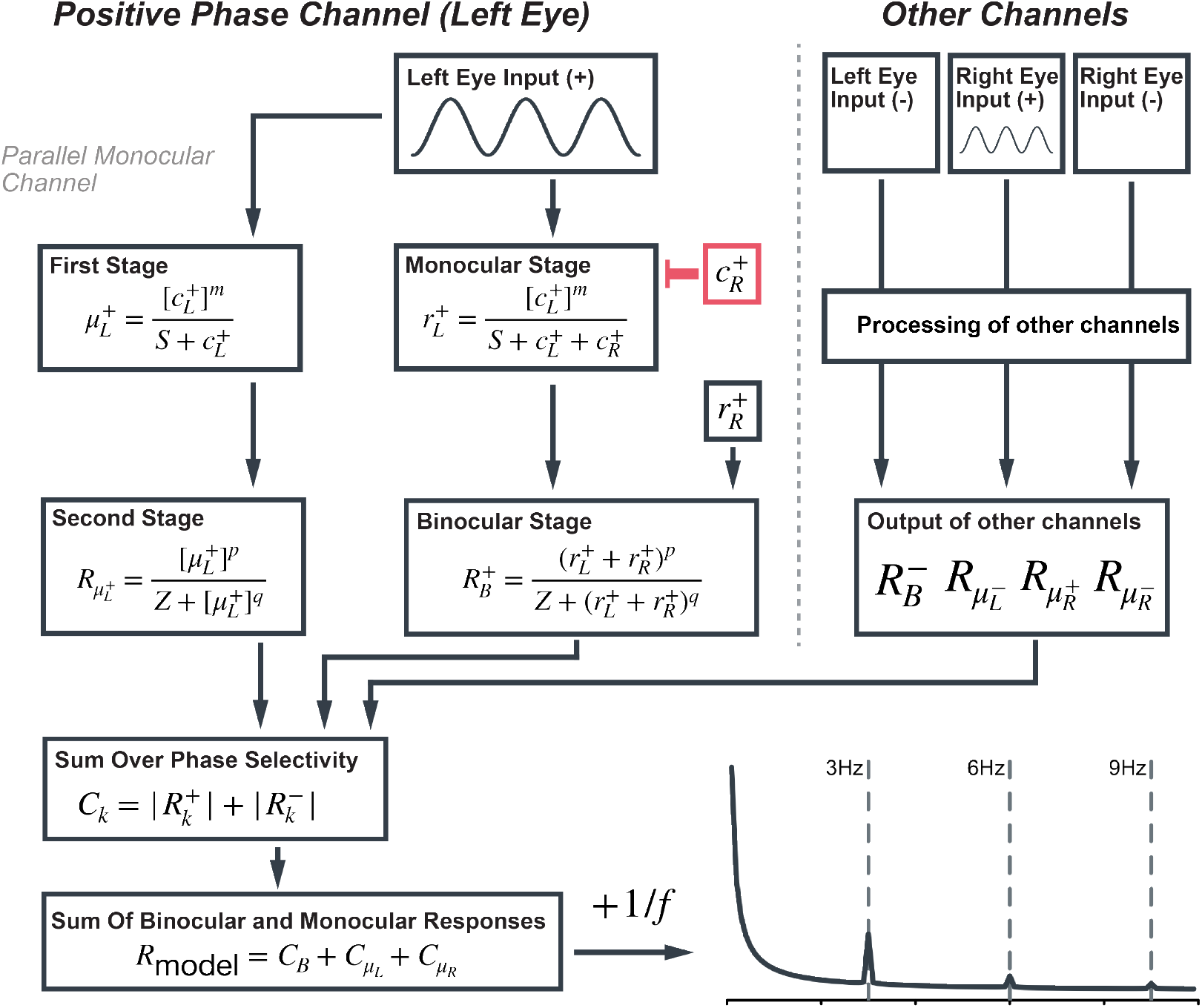
A simplified diagram of the two-stage contrast gain control model with parallel monocular and phase-selective channels. This diagram represents the processing stages of the positive phase channel in the left eye. These operations are identical for the other channels (negative phase left eye, positive and negative phase right eye) of the model. The sinusoidal input to the left eye 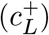 is fed through the first stage of the parallel monocular channel and the monocular stage of the binocular channel, which both apply a non-linearity to the input (*m*) and divisive inhibition (i.e., contrast gain control). The monocular stage of the binocular channel receives suppression from itself and the positive phase channel from the other eye. The output of these stages is fed into a second contrast gain control stage. It is at this stage that the monocular inputs are combined in the binocular channel. Finally, all responses are fast Fourier Transformed and their absolute values are summed over phase selectivity, followed by a sum over ocularity (binocular and monocular responses). This approach to defining the model’s output differs from that of Georgeson (2016) to account for methodological differences. A pink noise spectrum was added to facilitate a comparable calculation of model SNRs, as is done with human data.

The architecture of the models explored differs, but all received the same input and had their final outputs processed identically. The input to all models was a 3 Hz sine wave, adjusted to accurately represent the various experimental conditions of this study (see Figure 1). For example, stimuli presented with an On/Off flicker in temporal anti-phase had the left eye input generated by the following equation,

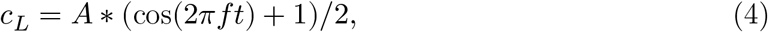

while the right eye input is defined as,

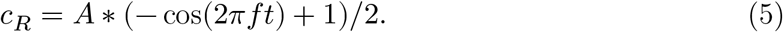

*A* represents stimulus contrast (amplitude), *f* the temporal flicker frequency (i.e., 3 Hz), and *t* time in milliseconds. The input to the right eye (*c*_*R*_) is phase shifted by 180°, which is accomplished using the negative cosine function (−*c*°*s*). Finally, sine waves are rectified to the range between 0 and 1 to represent the relative contrast presented to observers. The same experimental condition with counterphase flicker has the following sinusoidal profile for the left eye,

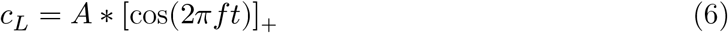

and for the right eye,

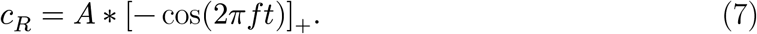

These profiles are identical to the On/Off flicker, but the sine waves are half-wave rectified to represent the counterphase oscillation. To fit model outputs (rectified sine waves) to observer data, the final response of the models was Fast Fourier Transformed, and a pink noise spectrum was added to the Fourier amplitudes, |*FFT*(*R*_model_)|+1/*f* to simulate neural noise (see e.g. Donoghue et al., 2020), before calculating model SNRs. All models developed in this study were fit by minimizing the sum of squared errors between the model output and the observer median SNRs for the first 6 SSVEP components (3Hz, 6Hz, 9Hz, 12Hz, 15Hz, and 18Hz).

### Evidently wrong models

As a first step in defining the necessary architecture to capture our results, we built baseline models with no monocular stage or phase selectivity that we do not expect to explain all of our effects. The first is a purely linear summation model of binocular combination;

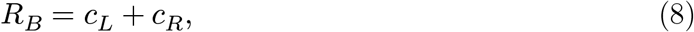

the binocular response (*R*_*B*_) is the sum of the inputs *c*_*L*_ and *c*_*R*_. The fits of the linear summation model are shown in Figure 5 and its performance metrics in Table 1. For On/Off flicker, the linear summation model only generates responses at the fundamental frequency (3Hz) that grossly overestimate observer SNRs. This is expected as this model lacks the rectification and non-linearities required to generate responses at the harmonics (Regan and Regan, 1988; Wade and Baker, 2025). In a linear sum, stimuli presented under On/Off flicker in temporal anti-phase cancel each other, and thus the model generates no response. The model does generate responses at the fundamental and harmonics of the counterphase flicker condition (Figure 5B), but this is attributable to the input’s rectification (the half-wave rectification applied to the input; Equation 6, Equation 7) and not the model architecture.

**Figure 5:**
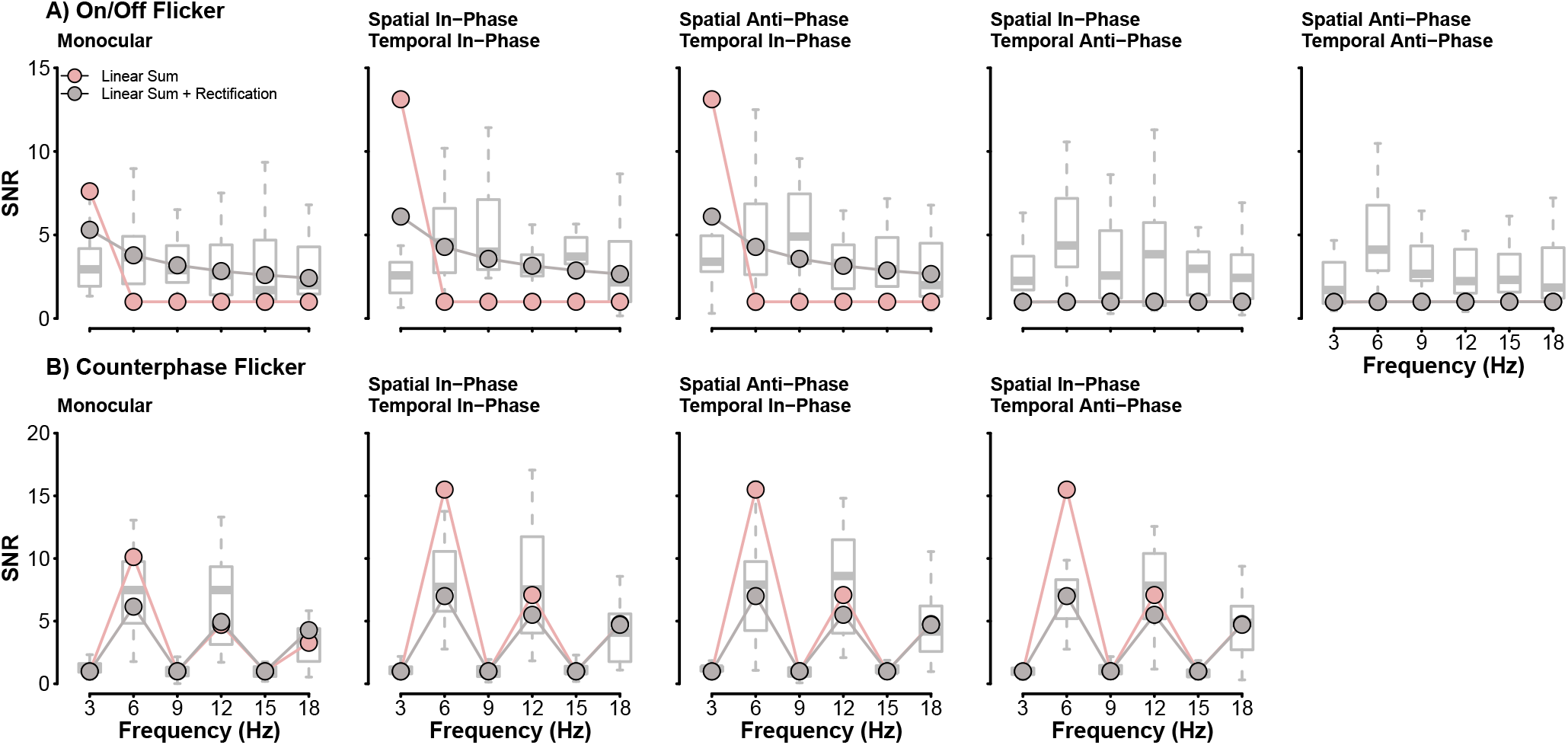
Fits of the linear sum (green) and the rectified linear sum (brown) models. Boxplots behind model responses show the distribution of observer SNRs. Model SNRs were fit to the median SNR of observers, which is represented by the thicker line within each box.

**Table 1:** Goodness-of-fit metrics for all models compared in this study. Errors in predictions for the linear sum model were too large to calculate *R*^2^. RMSE is the Root Mean Square error and AIC is the Aikaike Information Criterion. Note that, as the residuals of our fitting procedure are, by nature, skewed, the AIC values returned should be used only for relative comparisons of the models reported in this manuscript.

| Model | $R^2$ | RMSE | AIC |
| --- | --- | --- | --- |
| Linear sum | - | 3.234 | 281.99 |
| Linear sum, with Rectification | 0.575 | 1.405 | 197.95 |
| Two-Stage, no interocular interactions | 0.822 | 0.908 | 154.86 |
| Two-Stage, with interocular interactions | 0.81 | 0.94 | 158.62 |
| Two-Stage with parallel monocular channels | 0.877 | 0.756 | 135.1 |
| Two-Stage with phase-selective channels & monocular channels | 0.876 | 0.759 | 135.39 |

Responses of neurons to contrast in the visual system are well-modelled by a saturating non-linearity: as contrast increases, the magnitude of responses saturate (the rise in response per unit contrast decreases at higher contrast values (Heeger, 1992)). The saturating non-linearity can be modelled in different ways, but generally contains a divisive suppression and an exponentiation of the excitatory and inhibitory inputs. Including suppression can aid the model in better capturing the magnitude of responses in our observers, while exponents introduce the non-linearities required to generate responses at the harmonic frequencies. Thus, the next increment of model complexity defines the binocular response as the contrast gain control equation (after Legge, 1984b):

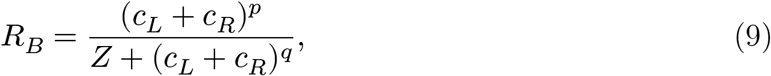

where the binocular response of the model (*R*_*B*_) is defined as the sum of monocular inputs (*c*_*L*_ and *c*_*R*_) raised to the power *p*, normalized by the sum of monocular inputs raised to *q* and where (*p* > *q*). The parameter Z sets gain of the model. The model can now generate responses at the harmonic frequencies for stimuli presented in temporal phase under On/Off flicker (see Figure 5 and Table 1). While this model iteration improves on the fits, it nevertheless struggles to fit SNR values at the fundamental frequency (3Hz) and is, as with the linear summation model, incapable of generating responses to stimuli presented with On/Off flicker in temporal anti-phase; the linear sum of stimuli presented in temporal anti-phase will always return zero. Therefore, the model still lacks the necessary architecture to define neural responses to our stimuli adequately.

### The Two-Stage Contrast Gain Control Model

The simple models described above could not accurately represent the observer SSVEPs we recorded. They overestimated SNRs at the fundamental frequency and failed to generate responses for stimuli presented in temporal anti-phase with On/Off flicker. A potential model refinement is adding a monocular transducer before binocular combination (Meese et al., 2006). The architecture of this model now begins with a monocular stage

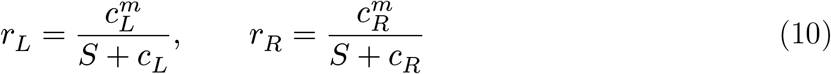

where an exponent (*m*) is applied to the monocular inputs (*c*_*L*_ and *c*_*R*_) in addition to self-suppression. The parameter *S*, as the parameter *Z* above, sets the overall sensitivity of the model. The outputs of the monocular stage (*r*_*L*_ and *r*_*R*_) are then fed into the binocular stage as defined in Equation 9,

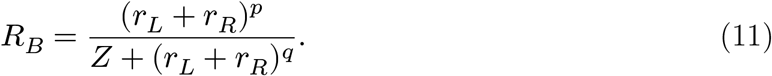

In this model variant, *m* is the monocular exponent that determines the extent of summation at detection threshold, with moderation by the suppressive term. In the second stage, *p* > *q* as with Equation 9, which is necessary to capture the mild facilitatory effects of dichoptic masking (Meese et al., 2006), and determines the shape of contrast discrimination functions. We can strengthen the normalization of the monocular input by adding interocular suppression and replacing Equation 10 (the first stage) with:

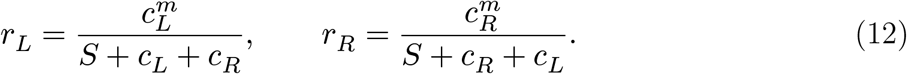

This model iteration is identical to the two-stage contrast gain control model defined by Meese et al. (2006).

The fits of both model variants, with and without interocular suppression, are shown in Figure 6. The difference in their quality is negligible (see Table 1) as both describe most experimental conditions well. The addition of the monocular transducer enables the model to fit observer SNRs at the fundamental frequency for stimuli presented in temporal phase under On/Off flicker, and importantly, it now generates responses for stimuli presented in temporal anti-phase. The transduced monocular inputs no longer cancel each other at the binocular stage. While the model can create responses to temporal anti-phase stimuli, it only does so at the even harmonics (2F-6Hz, 4F-12Hz, and 6F-18Hz) of the SSVEP spectrum. This is expected as the model can only generate binocular responses; it does not preserve monocular signals beyond the first stage. As the two rectified sine waves are in anti-phase, their sum will generate a new waveform with frequencies twice the original (6Hz) frequency and its integer harmonics (2F-12Hz and 3F-18Hz). The responses we recorded at the fundamental frequency (3Hz) and its odd integer harmonics (3F - 9Hz, 5F - 15Hz) cannot be explained by an architecture with a purely binocular output. Next, we explore methods of preserving the monocular signal to explain observer responses to stimuli presented in temporal anti-phase.

**Figure 6:**
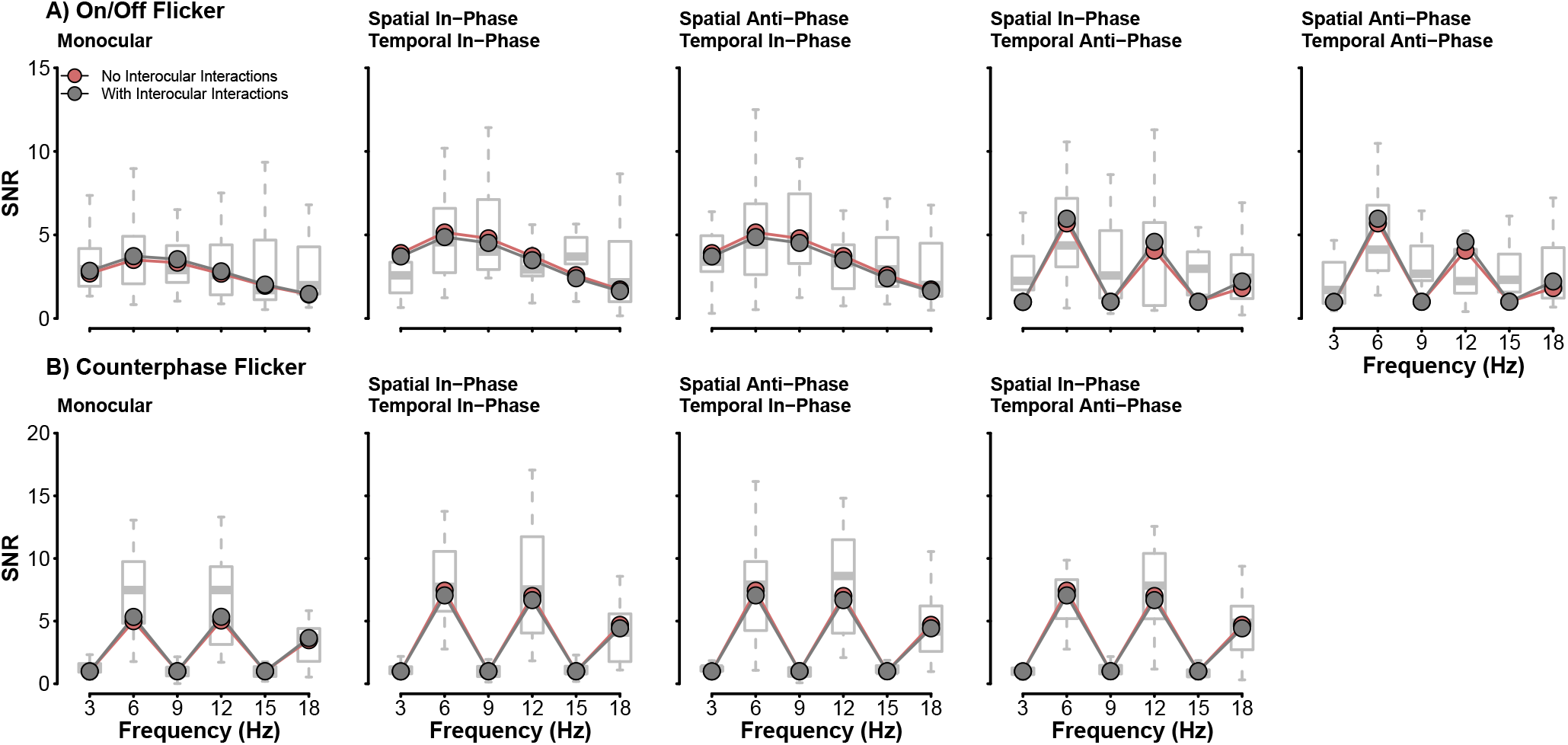
Fits of the two-stage contrast gain control model without (green) and with (red) interocular suppression. Boxplots behind model responses show the distribution of observer SNRs. Model SNRs were fit to the median SNR of observers, which is represented by the thicker line within each box.

### Parallel Monocular and Phase-Selective Channels

Based on the models described above, observer SSVEPs for stimuli presented in temporal anti-phase imply the presence of both a binocular and monocular response in the population response. To preserve the monocular response in the modelling, we add parallel monocular channels to the two-stage contrast gain control model, similar to Georgeson et al. (2016). These channels operate in an identical manner to the traditional two-stage contrast gain control model channel but are fully monocular (i.e., they do not include interocular suppression),

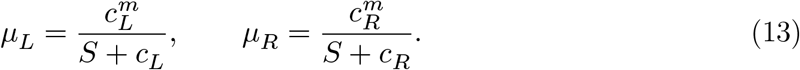

These equations, identical to those presented in Equation 10, return *μ*_*L*_, the output of the left eye’s first stage of the monocular channel, and *μ*_*R*_ is that of the right eye. The output of the monocular channels undergoes a second contrast gain control stage similar to that of the binocular channel,

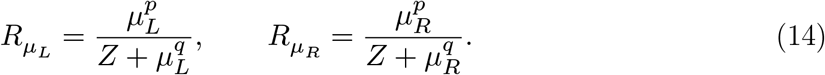

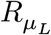 and 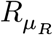 represent the final responses of the left and right monocular channels. No additional free parameters are required to define the parallel monocular channels; *m, p, q, S*, and *Z* are used to define the monocular channel and binocular channel responses.

The inclusion of parallel monocular channels poses an interesting problem in our modelling as we now contend with three visual cues from which to generate model SSVEPs: monocular left (*R*_*L*_), monocular right (*R*_*R*_), and binocular (*R*_*B*_). Psychophysically, cue selection has been implemented as a Minkowski sum with a large (≈ 30) exponent (Georgeson et al., 2016), approximating a MAX rule. This method is inappropriate when modelling neural data, as the recordings from the scalp represent an amalgamation of all responses generated by the stimuli (Wade and Baker, 2025). Our model output must therefore represent the combination of signals instead of the selection of the strongest signal. Here, it is implemented as the sum of the Fourier amplitude spectra of all three channel outputs. This sum preserves the amplitude responses required to generate SSVEPs and prevents the nullifying of responses from summing signals in anti-phase.

Preserving monocular responses until the output stage with parallel monocular channels improved the fit to our data (see Table 1 and Figure 7). The model now captures responses at the fundamental (3Hz) and odd-integer harmonics (9Hz and 15Hz) of the temporal anti-phase conditions under On/Off flicker. Monocular signals, while weaker than binocular signals, are preserved in the neural responses of observers and thus must be accounted for in their computational description. As proposed by Georgeson et al. (2016), parallel monocular channels appear to be an adequate descriptor of monocular signals.

**Figure 7:**
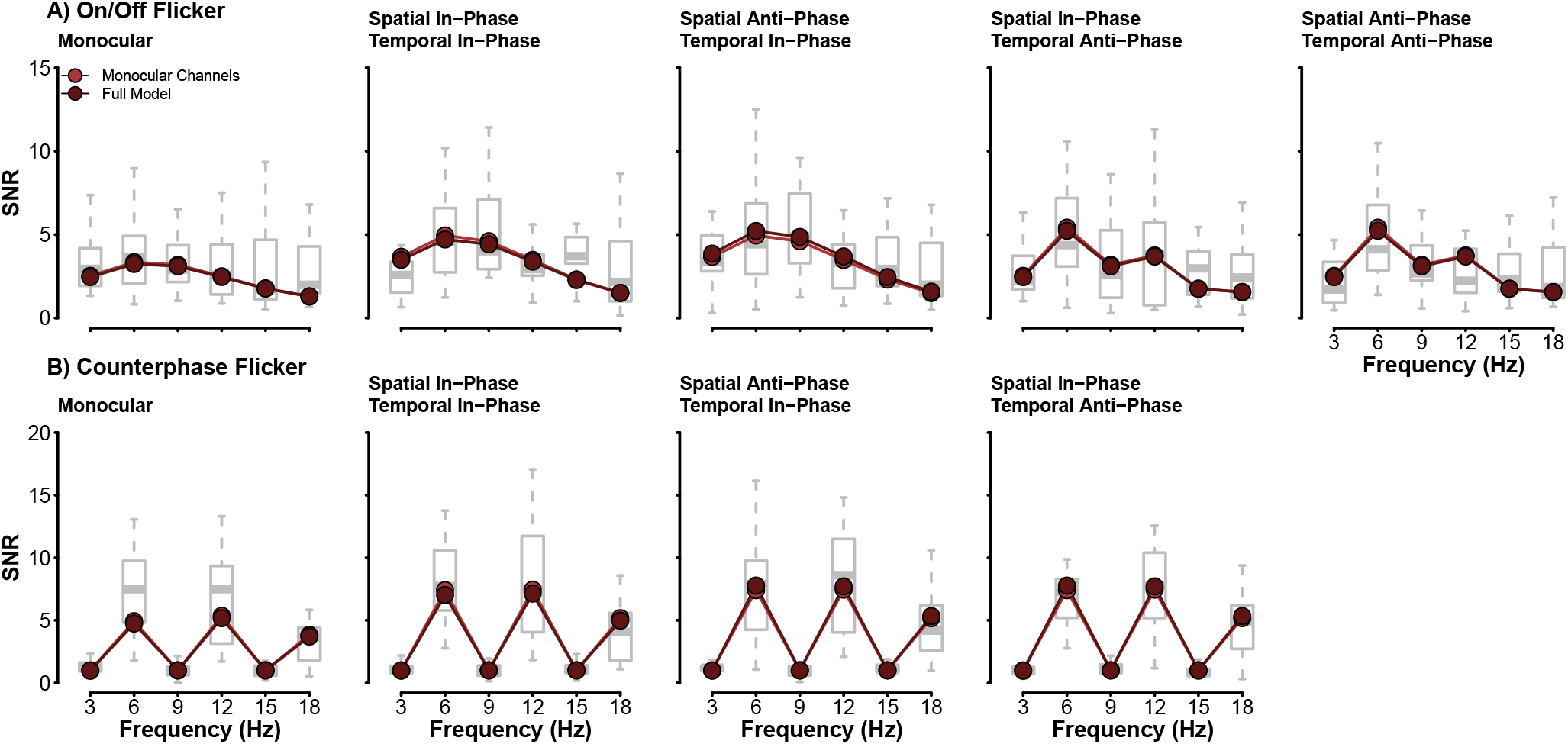
Fits of the two best performing models to our observer data. Both models, that including parallel monocular channels and the full model with the added phase-selective channels perform quite similarly.

Psychophysically, spatial phase has a meaningful impact on binocular combination (Bacon, 1976; Baker and Meese, 2007; Simmons, 2005) and the two-stage contrast gain control often includes phase-selective channels to account for these effects. Our study found little evidence of any influence of spatial phase on observer SSVEPs, as no statistically significant difference in signal-to-noise ratios across spatial phase was found. Still, we felt it prudent to verify if adding phase-selective channels improves the fit of the model to our data. Phase-selectivity was added to the binocular and monocular channels by replicating the equations of the first and second stage, for six channels (see Figure 4). As with the previous model, no additional model parameters were required to include phase-selectivity; the parameters *m, p, q, S*, and *Z* were used to define the responses of all six channels in this model.

Model responses were generated from sine waves (in temporal phase or anti-phase) fed into their respective positive or negative phase channels. This simulated the spatial phase of stimuli presented to observers. We refer to this model iteration as the full model, as it includes a binocular channel with a monocular stage (with interocular interactions), a binocular stage, parallel monocular channels (with their respective stages), and phase-selectivity. This is, in essence, the 6-channel 2-cue model proposed by Georgeson et al. (2016) to explain binocular combination under multiple experimental conditions. As with the addition of monocular channels, including phase-selectivity to the model means we now have six final signals to contend with: a binocular and two monocular channels selective for positive spatial phase, and a binocular and two monocular channels selective for negative spatial phase. We used the same method of combining signals as above: we first summed the Fourier amplitude spectra across the positive and negative phase-selective channels and then combined the binocular and monocular channels. The inclusion of phase-selectivity did not improve the model’s fit. The *R*^2^ value calculated across the six frequencies and nine experimental conditions was no different than that of the previous model (see Table 1), indicating that the spatial phase of stimuli does not have a meaningful impact on the SSVEP amplitudes of observers.

## Discussion

The computational architecture of binocular combination, under the two-stage contrast gain control model framework, has been carefully evaluated on psychophysical data (Baker and Meese, 2007; Georgeson et al., 2016; Meese et al., 2006), yet its ability to explain neural data has only been explored on a limited set of stimulus conditions (Baker and Wade, 2017; Lygo et al., 2021; Richard et al., 2018). This study investigated the effects of stimulus spatial and temporal phase on observer SSVEPs in an effort to probe the ability of the two-stage contrast gain control model - and its variants - to capture our neural data. On/Off flicker generated responses at the fundamental frequency (3Hz) and its harmonics (6Hz, 9Hz, 12Hz, and 15Hz) while counterphase flicker generated responses at twice the fundamental frequency (6Hz) and its even integer harmonics (12Hz and 18Hz). No statistically significant difference was found between the monocular and binocular conditions presented with On/Off or counterphase flicker. This is consistent with ocularity invariance; binocular and monocular stimuli appear equal in magnitude at high contrast (Baker et al., 2007a; Meese et al., 2006). We additionally found no statistically significant differences in SSVEPs for all stimulus conditions under counterphase flicker.

Our data proved useful in selecting the best-performing model, albeit in an unexpected way. The stimulus presentation protocols (On/Off and counterphase) individually were unable to discern the best model, as all models, except the evidently wrong ones, were able to capture neural responses to counterphase stimuli. Instead, it was the spatial and temporal phase of our stimuli that proved most useful in model selection. Presenting stimuli in temporal antiphase reduced the magnitude of responses at the fundamental frequency (3Hz) compared to its temporal phase counterpart, but they were not abolished. This finding indicates that, even under binocular viewing, monocular signals can be measured at the scalp with SSVEPs. This was crucial in the selection of the best fitting model. Only the two-stage contrast gain control model with parallel monocular channels, which preserves monocular signals throughout the model architecture, could accurately capture the data in all experimental conditions. Simpler models failed to generate the necessary responses at the fundamental and odd-integer harmonics for stimuli presented in temporal anti-phase. In contrast, a more complex model that included phase selectivity did not improve the quality of fits. Overall, we find that the same general framework used to explain psychophysical stimulus combination can be successfully applied to neural data collected under various experimental conditions.

### Monocular channels

It has long been assumed by most computational descriptions of binocular combination that only the binocular response was available to later stages of perception and decision (Baker and Meese, 2007; Ding et al., 2013; Ding and Sperling, 2006; Legge, 1984a; Meese et al., 2006). The monocular pathways, which represent the early stages of the visual processing stream, only serve as the input to combine. However, psychophysical evidence from adaptation and discrimination experiments has demonstrated that monocular signals do contribute to perception (Blake et al., 1981a; Georgeson et al., 2016; Moulden, 1980) and that monocular channels parallel to the binocular channel should be included in models of binocular combination. This study found that monocular responses to stimuli can be recorded with SSVEPs under binocular stimulation. Presenting stimuli in temporal anti-phase under On/Off flicker generates two transients per cycle, and a binocular system, which combines the monocular inputs of each eye, would only create responses at 6Hz. While the 6Hz component was significant, we still found SSVEPs at 3Hz that exceeded the noise level in our data. These 3Hz components represent the individual oscillations for each eye and are therefore likely representative of a monocular response. The reduction in response magnitude of the 3Hz component we observed from the temporal in-phase to the temporal anti-phase conditions can thus be explained as the transition from a binocular to an inadvertently weaker monocular response.

Computationally, we demonstrated that many mechanisms of binocular combination, including monocular non-linearities, interocular interactions, and parallel monocular channels, were required to explain the SNR spectra of our observers. Parallel monocular channels were critical in capturing observer data in the two temporal anti-phase conditions presented with On/Off flicker. Without adding these channels, the models could not generate a response at the fundamental (3Hz) or its odd integer harmonics. While an essential inclusion to explain our data, the addition of monocular channels did not meaningfully alter the ability of the two-stage contrast gain control model to capture observer SNRs in the other experimental conditions. Monocular contributions to the model output are most significant for perception when the stimulus contrast between the two eyes differs (Georgeson et al., 2016), an experimental scenario we did not explore here.

### SSVEP responses to spatial phase

Our results did not suggest an impact of stimulus spatial phase on observer SSVEPs. It is well-known that spatial phase affects binocular combination when measured psychophysically (Bacon, 1976; Baker and Meese, 2007; Simmons, 2005). The binocular combination of two opposite-polarity stimuli does not cancel; it sums, thus influencing sensitivity to the stimuli. It is therefore odd to find little to no influence of spatial phase on the SSVEPs of observers, particularly as SSVEP amplitude is associated with perceptual sensitivity (Bosse et al., 2018; Campbell and Kulikowski, 1972; Campbell and Maffei, 1970; Norcia et al., 2015). However, we note that at high contrasts (as used here), perceived contrast is equivalent between inphase and antiphase stimuli (Baker et al., 2012). The absence of an apparent spatial phase effect also complicates our ability to investigate any contributions of a differencing channel in our study (Chen and Li, 1998; May et al., 2012; May and Zhaoping, 2016).

Spatial phase-dependent effects in SSVEPs have been recorded with motion stimuli (Cottereau et al., 2014; Kohler et al., 2018). The dichoptic presentation of moving bars, whereby they either move in-phase (lateral motion) or in anti-phase (motion-in-depth), impacts the amplitude of SSVEPs. In-phase motion generates SSVEP amplitudes that are twice those of motion in anti-phase (Cottereau et al., 2014). A different response can also be recorded for dichoptic stimuli when frequency tagged (Katyal et al., 2018; Sutoyo and Srinivasan, 2009), where the “conflict” response is represented at the intermodulation frequencies. All stimuli in this study were presented at 3Hz; we did not frequency tag stimuli presented to the left and right eyes. Thus, the responses we recorded at the scalp are a spatial aggregation of the stimuli presented to observers (Wade and Baker, 2025). Spatially aggregated responses will be identical regardless of stimulus phase and thus generate phase-insensitive responses.

## Conclusion

We investigated the effects of stimulus spatial and temporal phase on observer SSVEPs and explored the necessary computational components to explain our data. We worked under the framework of the two-stage contrast gain control model of binocular vision (Meese et al., 2006). We examined the impact of interocular interactions, parallel monocular channels, and phase-selective channels on the model’s fit with our data. The most significant effect to capture was the presence of odd-integer harmonics in the temporal anti-phase conditions, representing the monocular response to stimuli. This was well-explained by adding parallel monocular channels to the model. The two-stage contrast gain control model of binocular combination remains a robust descriptor of binocular vision as it can explain various experimental conditions and modalities (psychophysics and neuroimaging).

## Data and Code Availability

We provide the raw (.cnt) and processed (.RData) observer SSVEPs, in addition to all code used in this study, on the Open Science Framework, which can be accessed using the following project link: https://osf.io/cn894/. A computationally reproducible version of this manuscript is available at the linked GitHub repository:https://github.com/brunoRichard/SSVEP_Phase_AntiPhase.

